# HPV-positive head and neck cancer derived exosomal miR-9 induces M1 macrophage polarization and increases tumor radiosensitivity

**DOI:** 10.1101/820282

**Authors:** Fangjia Tong, Siwei Zhang, Huanhuan Xie, Bingqing Yan, Lianhao Song, Lanlan Wei

**Affiliations:** Department of Microbiology, Harbin Medical University, 157 Baojian Road, Harbin 150081, China; Wu Lien-Teh institute, Harbin Medical University, 157 Baojian Road, Harbin 150081, China

**Keywords:** HPV, Exosome, miR-9, Radiotherapy, Head and neck cancer

## Abstract

Human papillomavirus (HPV) is an etiological risk factor for a subset of head and neck squamous cell carcinoma (HNSCC). HPV+ HNSCC is significant more radiosensitive than HPV-HNSCC, but the underlying mechanism is still unknown. Tumor microenvironment can affect tumor response to radiation therapy. Cancer secreted exosomes are emerging as crosstalk mediators between tumor cells and the tumor microenvironment. The main objectives of this study were to determine the role of HPV+ HNSCC-derived exosomes in increased radiation sensitivity. Here, we found that exosomes derived from HPV+ HNSCC cells activate macrophages into the M1 phenotype, which then increases the radiosensitivity of HNSCC cells. miR-9 was enriched in exosomes released from HPV+ HNSCC cells and it could be transported to macrophages, leading to altered cellular functions. Overexpression of miR-9 in macrophages induced polarization into the M1 phenotype via downregulation of PPARδ. Increased radiosensitivity was observed for HNSCC cells co-cultured with macrophages in which miR-9 was upregulated or treated with M1 macrophages. These observations suggest that HPV+ HNSCC cells secrete miR-9-rich exosomes, which then polarize macrophages into M1 phenotype and lead to increased radiosensitivity of HNSCC cells. Hence, miR-9 may be a potential treatment strategy for HNSCC.

**Statement of significance:** HPV+ HNSCC through the release of miR-9-rich exosomes polarize macrophages into M1 phenotype and lead to increased radiosensitivity of HNSCC.

## Introduction

Head and neck squamous cell carcinoma (HNSCC) is the sixth most common cancer worldwide with an annual incidence of approximately 600,000 cases (1,2). HPV has been considered to play an etiological role in HNSCC oncogenesis, especially HPV-16 (3). Depending on the anatomic site of the tumor, HPV prevalence in HNSCC ranges from 10% to 90% (4,5). HPV+ HNSCCs show a distinct subset of HNSCCs that differs from HPV-HNSCCs in tumor biology and clinical characteristics, including improved radiosensitivity (6).

Over the last few years, many papers have suggested that HPV+ HNSCCs had a better response to ionizing radiation (IR). There are many factors that contribute to improved radiosensitivity, including the presence of wild-type p53, lower levels of tumor hypoxia, tumor immunology, and so on. However, the potential underlying mechanisms are still unknown. Immune cells within the tumor microenvironment (TME) are reported to be associated with tumor radiosensitivity (7,8). As lymph nodes and vessels are abundant in the head and neck region, tumor immunity may play a role in enhancing the tumor response to IR (9). Macrophages, the most abundant infiltrative immune-related stromal cells present in and around tumors, display diverse phenotypes and functions (10,11). Macrophages can be polarized into classical M1 or alternatively activated M2 macrophages, depending on the environmental cues (12–14). In a previous study, we found that HPV+ HNSCCs contained relatively more M1 macrophages, while HPV-HNSCCs had more M2 macrophages. The ratio of M1/M2 was significantly increased in HPV+ HNSCCs (15).

Exosomes are tiny vesicles 30 to 150 nm in size that are formed during endocytosis (16). It has been reported that exosomes transfer their contents, including mRNAs, miRNAs and proteins, to immune cells and play an essential role in the communication between tumor and the surrounding environment (17). Recently, several studies have reported that exosomes derived from tumor cells play an important role in immune modulation. Ye SB et al. showed that exosomal miR-24-3p impedes T-cell function (18). Xiang Y et al. showed that epithelial ovarian cancer-secreted exosomal miR-222-3p induced polarization of tumor-associated macrophages(19).

In this study, we demonstrate that HPV+ HNSCC-derived exosomal miR-9 can polarize macrophages into the M1 phenotype. Polarization into M1 macrophages leads to improved radiosensitivity of HNSCC cells. Thus, the data implicate miR-9 as a potential aid for the treatment of HNSCC.

## Materials and Methods

### Patient samples

A total of 47 HNSCC samples were collected from the Third Affiliated Hospital of Harbin Medical University (Harbin, China), according to the Ethics Committee of Harbin Medical University. Informed consent was obtained from patients or their guardians. Tissue samples were paraffin embedded and rapidly frozen in liquid nitrogen for subsequent analysis. All samples were obtained from patients who were not treated with radiation or chemotherapy before surgery.

### Cell culture and treatment

Human HNSCC cell line SCC90 was purchased from American Type Culture Collection (Rockville, MD). SCC47, SCC104, SAS, CAL33 cell lines were kindly provided by Dr. Henning Willers (Harvard University, Boston, USA). HPV-cell line CAL27 was a kind gift from Professor Songbin Fu (Harbin Medical University, Harbin, China). HNSCC cell lines were maintained at 37°C in a humidified atmosphere of 5% CO2 and cultured in DMEM supplemented with 10% exosome-free fetal bovine serum (FBS). To obtain exosome-free FBS, FBS was ultra-centrifuged at 120,000 × *g* for 20h.

### Macrophage differentiation and polarization

Human monocytic cell line THP1 was obtained from American Type Culture Collection (Rockville, MD) and maintained in RPMI 1640 medium containing 10% FBS. To obtain resting macrophages (M0), THP1 cells (1×10^6^) were treated with 100ng/ml 12-myristate 13-acetate (PMA) (Sigma-Aldrich, St Louis, MO, USA) for 24h and rested for another 24h. M0 cells were polarized into M1 using 50 ng/ml IFN-γ (Beyotime, China) and 1μg/ml LPS (Beyotime, China) for 24h.

### Exosome isolation and identification

Collected cell culture medium was centrifuged at 300 × *g* for 10 min, 2000 × *g* for 30 min (to remove cells and debris), and ultra-centrifuged at 10,000 × *g* for 30 min. This was followed by ultra-centrifugation at 100,000 × *g* for 70 min to collect pellet that was resuspended in 50-100μl PBS. Exosomes to be examined by transmission electron microscopy (TEM) were dissolved in PBS buffer, dropped in a carbon-coated copper grid and then stained with 2% uranyl acetate. Samples were observed using a J Tecnai G2 F20 ST transmission electron microscope.

### Exosome labeling and tracking

Exosomes isolated from the culture medium of HNSCC cells were collected and stained with PKH67 Green Fluorescent membrane linker dye (Sigma-Aldrich), according to the manufacturer’s instructions. Labeled exosomes were resuspended and added to the unstained macrophages for exosomes uptake studies. After incubation for 2h at 37°C, cells were observed using a Leica DM-LB confocal microscope.

### Immunohistochemical Staining

Tumors were fixed, embedded in paraffin and sectioned at a thickness of 4 μm. After deparaffinization and rehydration, sections were blocked with 3% H2O2 and incubated with iNOS antibody (1:100) (Abcam, MA, USA), TNF-a antibody (1:200) (Abcam, MA, USA), CD163 antibody (1:200) (Abcam, MA, USA), CD9 antibody (1:100) (Proteintech, MA, USA), PPARδ antibody (1:200) (Proteintech, MA, USA) overnight at 4°C, followed by incubation in secondary biotinylated antibody for 30 min at 37°C. Blots were visualized with DAB solution and counterstained with haematoxylin. Pictures were taken under a light microscope. Stained slides were reviewed and scored independently by two observers blinded to the samples’ information. Scores were determined by a combination of the proportion of positively stained cells and the intensity of staining as follows: 0 (no positive cells), 1 (< 10% positive cells), 2 (10–50% positive cells), and 3 (> 50% positive cells). Staining intensity was classified according to the following criteria: 0 (no staining), 1 (weak staining = light yellow), 2 (moderate staining = yellow brown), and 3 (strong staining = brown). Staining index was calculated as the staining intensity score × the proportion score. Using this method, IHC data was evaluated by determining the staining index with scores ranging from 0, 1, 2, 3, to 9.

### Immunofluorescence-based radiosensitivity assay

Frozen tissue sections were fixed in 10% neutral formaldehyde for 15 min, permeabilized with 0.5% Triton X-100, and then blocked for 30 min with 5% BSA. Sections were incubated overnight at 4°C with mouse anti-γH2AX (Ser139) (Merck-Millipore, Germany). Following three washes with PBS, the sections were incubated with secondary antibody, washed three times and then mounted in anti-fade mounting medium containing DAPI (Solarbio, China).

### Western blotting

Western blotting was performed as previously described. Cells or exosomes were collected and denatured. Proteins were separated by SDS-PAGE gel and transferred onto polyvinylidene difluoride membranes. Following blocking in 5% skim milk for 30 minutes, membranes were probed with various primary antibodies overnight at 4°C, followed by incubation with horseradish peroxidase–linked secondary antibodies for 1h at room temperature. Visualisation was performed using electrochemiluminescence.

### ELISA assay

Cell culture medium of macrophages stimulated with HNSCC-derived exosomes or treated with miR-9 mimics and the miR-negative control was collected. Expression of following was measured using ELISA kits – IL-6, TNF-α and IL-10 (R&D System Co., Abington, UK).

### Real-time PCR

Total RNA was extracted from cultured cells using the mirVana kit (Invitrogen, USA) as per manufacturer’s instructions. Reverse transcription was performed using the High Capacity cDNA synthesis kit (Life Technologies, USA). Each RT reaction included total RNA as the template and a pool of miRNA-specific RT primers. Real-time PCR was performed individually for miRNAs with Power SYBR Green PCR Master Mix (Life Technologies, USA). GAPDH and U6 served as internal controls for expression data normalization. Expression changes were determined using the 2ΔΔCt method by comparison with the negative control.

### Luciferase assays

Reporter plasmid containing the 3′ untranslated region (3′ UTR) of the PPARδ gene was generated by cloning the 3′ UTR downstream of the luciferase open reading frame (Hanyin Biotechnology, Shanghai, China), The cloned plasmid together with the miR-9 mimics or the miR-negative control was transfected using Lipofectamine 2000 (Invitrogen, CA, USA) into HEK293T cells in a 24-well plate. 500ng of plasmid and 50 nM of miR-9 mimic were used for transfection. A constitutively expressed Renilla luciferase was co-transfected as a normalisation control. Following incubation for 40h, Firefly and Renilla luciferase activities were sequentially measured using the Dual-Glo Luciferase Assay system (Promega, Madison, WI, USA).

### Irradiation

All specimens were collected in tubes containing saline and brought into 37°C incubator within 1h (or so) of surgery. The tissues were cut using a scalpel into 3–5 mm pieces – one served as untreated control, and other one was irradiated. Tissues were cultured in 10% RPMI 1640 medium and received irradiation immediately in our lab. The optimal dose of X-ray irradiation was 6 Gy (RadSource RS2000, US). The specimens were cultured at 37°C in a humidified incubator for 24h. After 24h, the tissues were rinsed with fresh PBS and rapidly frozen in Optimal Cutting Temperature (OCT) compound (SAKURA, Japan) using liquid nitrogen. Tissues were cut with a frozen microtome while embedded in OCT. Tissue slices were placed on glass slides and stored at −80 °C in sealed slide boxes. Thickness of tissue slices was approx. 3–5 μm.

### Small RNA sequencing

Exosomes were isolated from cell culture medium as described above. Total exosomal RNA was extracted using the miRNeasy micro Kit (Qiagen, USA). Small RNA libraries were prepared using the NEBNext small RNA library preparation kit (New England Biolabs, USA). cDNA library was sequenced using Illumina Hiseq 3000 platform. Raw sequence reads were preprocessed using a custom bioinformatics pipeline for clustering before mapping to the RefSeq human miRNAs with Bowtie. Mapped sequence reads were then normalized using the RPKM method (reads per kb per million), and compared across samples to evaluate changes in expression.

### Bioinformatic analyses

Bioinformatic analyses were performed based on the TCGA HNSCC cohort using the UCSC Xena Browser. In total, 604 cases were searched for miRNA expression data. Only primary HNSCC cases were included for analysis.

### Statistical analysis

All statistical analyses in this study were conducted using SPSS16.0 software (SPSS, Inc., Chicago, IL, USA), and data are presented as mean ± SD. Significant difference between two groups was determined using Student’s t-test. Pearson Chi-squared tests were performed to assess the statistical significance for correlation between two variables. *P*<0.05 was considered as statistically significant.

## Results

### HPV promotes secretion of exosomes from HNSCC cells to induce macrophage M1 polarization

To examine the impact of HPV on exosomes released from HNSCC cells, exosomes of HPV+ and HPV- HNSCC cells were isolated from cell culture medium and quantitated by TEM. As shown in Figure 1A, TEM revealed the morphology and size of the acquired exosomes. The exosomes showed in Figure 1A are typical rounded particles ranging from 50 to 150nm in diameter. Moreover, exosomes harvested from HPV+ HNSCC cells demonstrated significantly higher nanoparticle concentrations compared with exosomes derived from HPV-HNSCC cells. Exosome identity was confirmed by the presence of known exosomal proteins – CD9, CD63 and TSG101. Increased levels of CD9, CD63, TSG101 were observed in the isolated exosomes (Figure 1B). Consistent with these observations, the expression level of CD9 (exosomes marker) examined by IHC in HPV+ HNSCC tissues was significantly higher than HPV- HNSCC tissues (Figure 1C). These results demonstrated that HPV enhances exosomes secretion in HNSCC cells.

**Figure 1.**
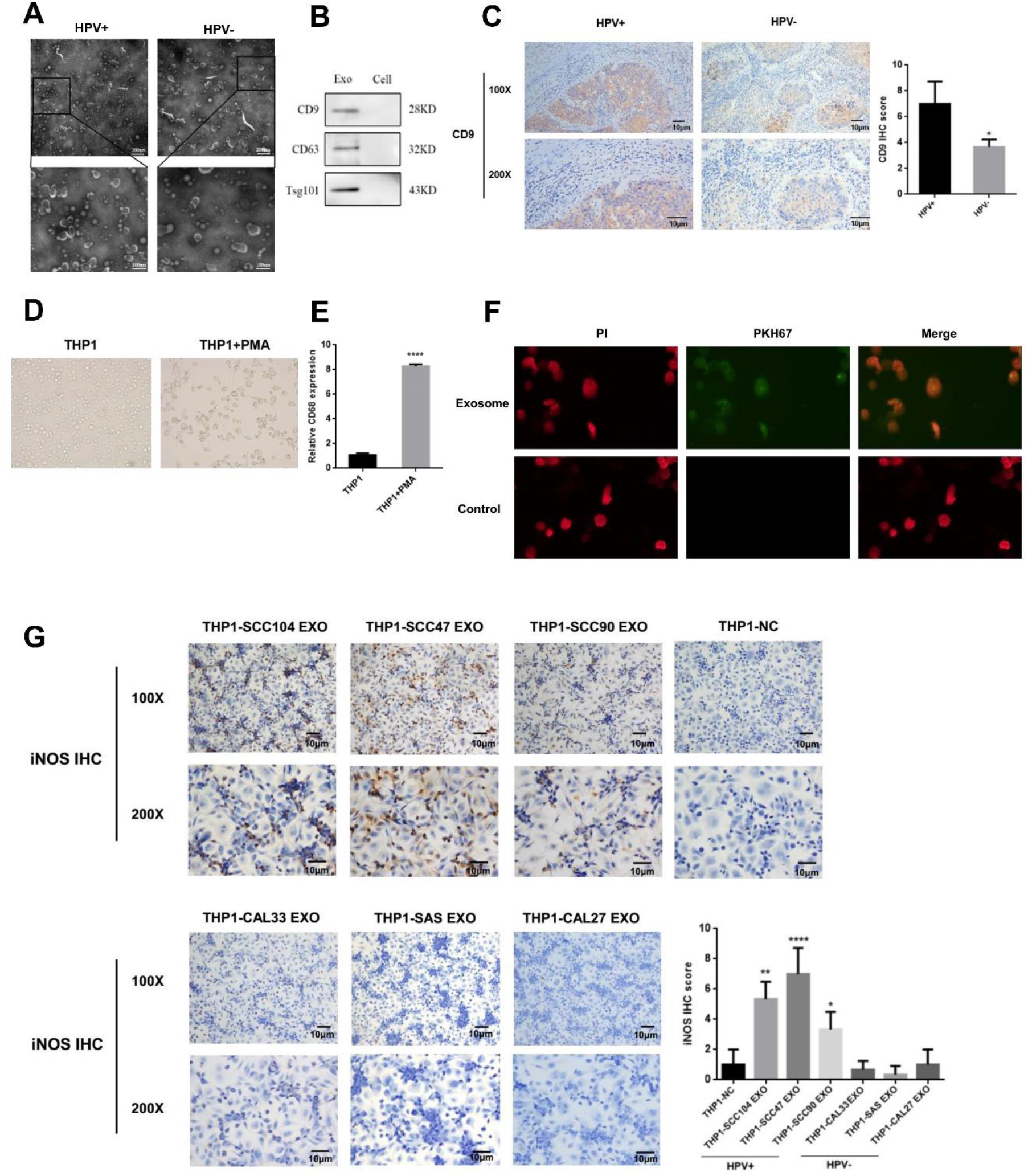
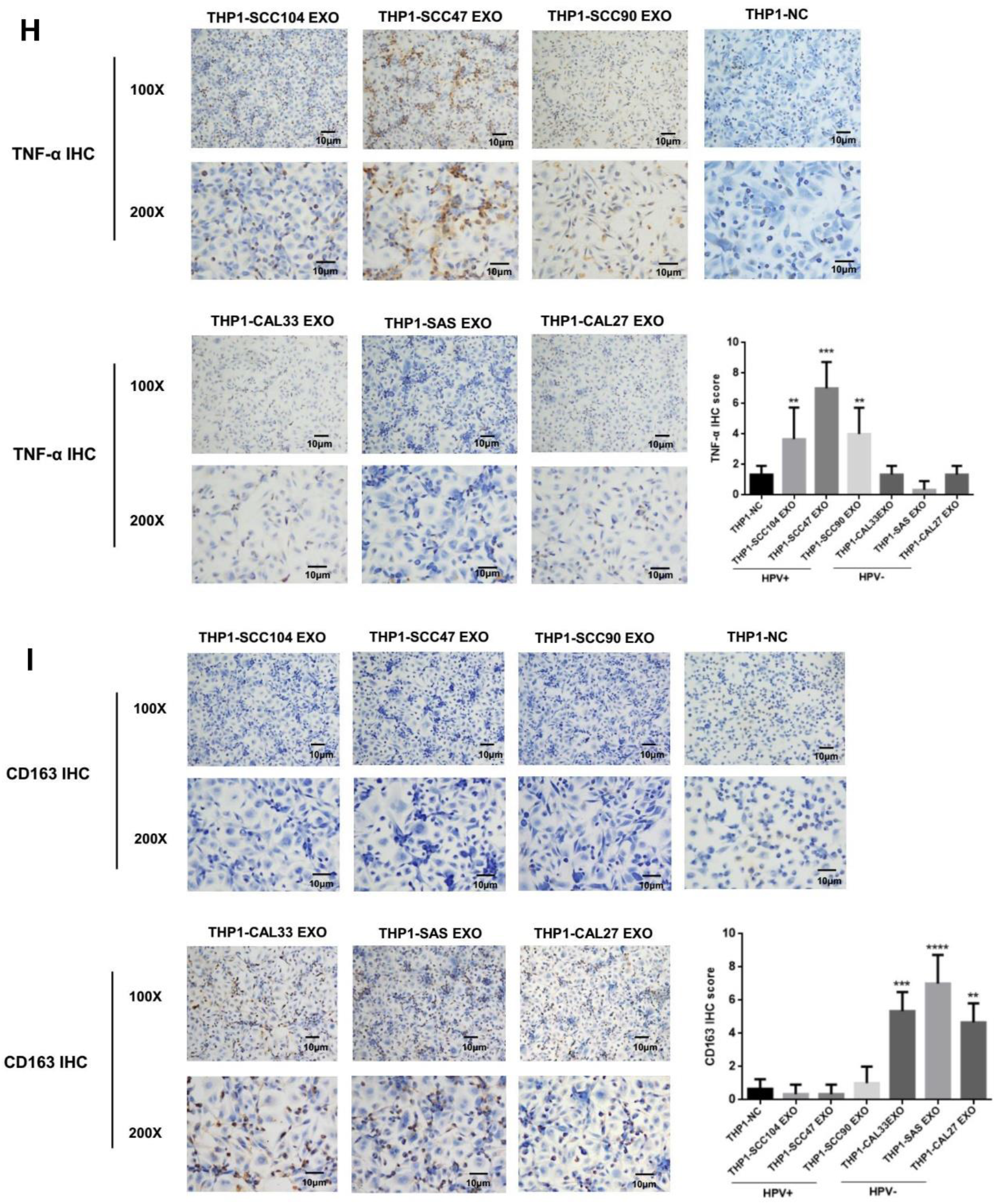
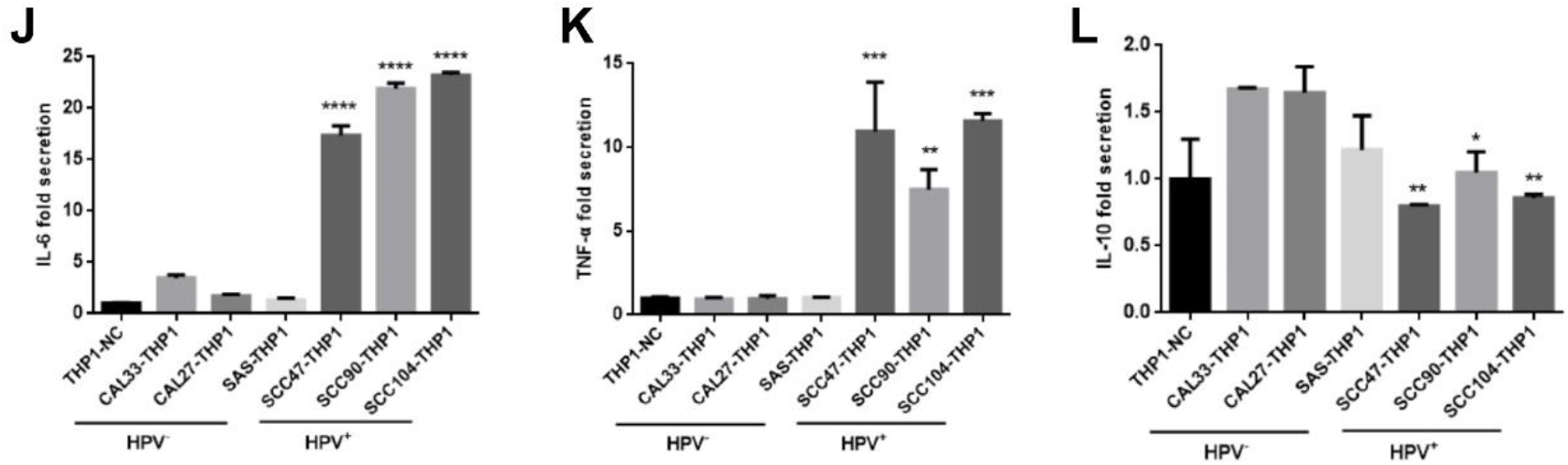
HPV promotes the secretion of exosomes from HNSCCs to induce the polarization of M1 macrophages. **A**, Representative TEM micrographs of exosomes derived from HPV+ and HPV-HNSCC cells. **B**, Western blot analysis for exosomal proteins – CD9, CD63 and TSG101. **C**, Expression of CD9 was examined by IHC in HNSCC. **D**, THP1 cells were treated with PMA for 24h. A representative image for macrophages is shown. **E**, qPCR was used to detect the expression of CD68. **F**, Immunofluorescence image shows the internalization of PKH67-labeled HNSCC-derived exosomes (green) by macrophages (red). Macrophages were treated with exosomes derived from HPV+ and HPV-HNSCC cells or control (PBS). After 48h incubation, IHC was used to detect the expression of M1 markers (iNOS, TNF-α) (**G-H**) and M2 marker (CD163) (**I**). **J-L**, Cytokine levels in culture medium of treated macrophages. \**P*<0.05, \*\**P*<0.01, \*\*\**P*<0.001, \*\*\*\**P*<0.0001.

HPV status has important implications for HNSCCs and secreted exosomes could deliver biological information by internalization into neighboring or distant cells. Macrophages are the most abundant infiltrative immune-related stromal cells present in and around tumors. Thus, we investigated whether exosomes derived from HPV+ HNSCC cells could affect the polarization of macrophages. Human monocytes THP1 were differentiated into macrophages by incubating in the presence of PMA, THP1 derived macrophages were characterized by its adherent morphology and the expression of recognized macrophage marker CD68 (Figure 1D-E). We next examined the effects of HPV+ HNSCCs derived exosomes on macrophages polarization. Exosomes labeled with fluorescent PKH67 were co-cultured with unstained macrophages. Examination with a confocal microscope revealed that exosomes were internalized by unstained macrophages over time (Figure 1F). IHC was used to comparatively analyze expression of macrophage subpopulation markers TNF-α, inducible nitric oxide synthase (iNOS, specific for M1 macrophages), and CD163 (M2 macrophages). The results showed the expression of M1 markers (iNOS and TNF-α) increased in macrophages treated with HPV+ HNSCC-derived exosomes compared with HPV-HNSCC-derived exosomes or PBS (Figure 1G-H), whereas HPV-HNSCC-derived exosomes increased expression of M2 marker CD163 (Figure 1I). M1 macrophages produce higher levels of IL-6 and TNF-α whereas M2 macrophages secret IL-10. Consistent with the changes in surface markers, macrophages treated with HPV+ HNSCC-derived exosomes produced a higher level of IL-6 and TNF-α, but a lower level of IL-10 (Figure 1J-L).

### M1 macrophages induced by HPV+ HNSCC-derived exosomes increase tumor radiosensitivity

We next investigate the role of M1 macrophages induced by HPV+ HNSCC-derived exosomes in the radiosensitivity of HNSCCs. As shown in Figure 2A, M1 macrophages were significantly more prevalent in HPV+ HNSCCs, while M2 macrophage were less abundant in HPV+ HNSCCs compared with HPV-HNSCCs (*P*<0.01). HPV+ and HPV-HNSCC tissue samples were treated with 6 Gy X-ray radiation. Within 24h of irradiation, the γ-H2AX foci were assessed by immunofluorescence (Figure 2B). Quantitative IF analysis showed that the radiation treatment increased the values of γ-H2AX foci for HPV+ HNSCC samples (*P*<0.001). The increase was significantly higher than observed for HPV-HNSCC samples (Figure 2C). Correlation analysis between M1 macrophage prevalence (based on IHC examination of markers) and radiosensitivity of HNSCC (based on γ-H2AX foci) were performed using Spearman model (Figure 2D). Our results indicated that M1 macrophages enhanced the radiosensitivity of HNSCC cells.

**Figure 2.**
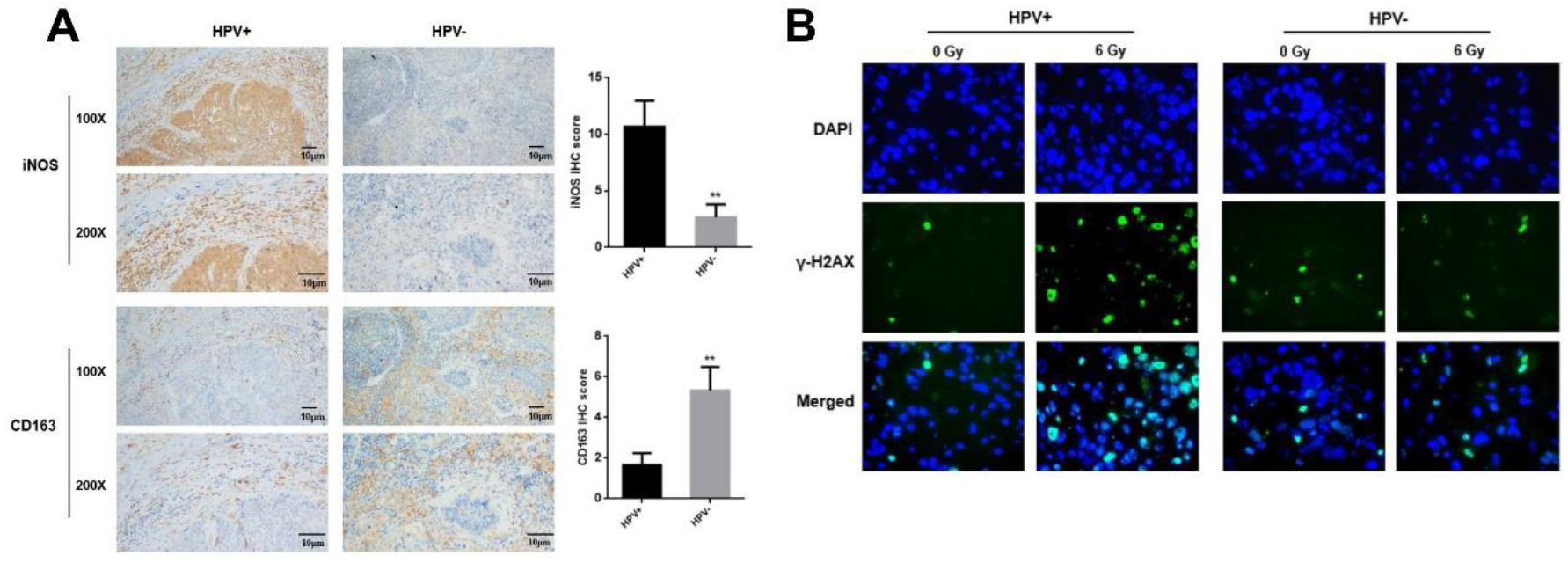
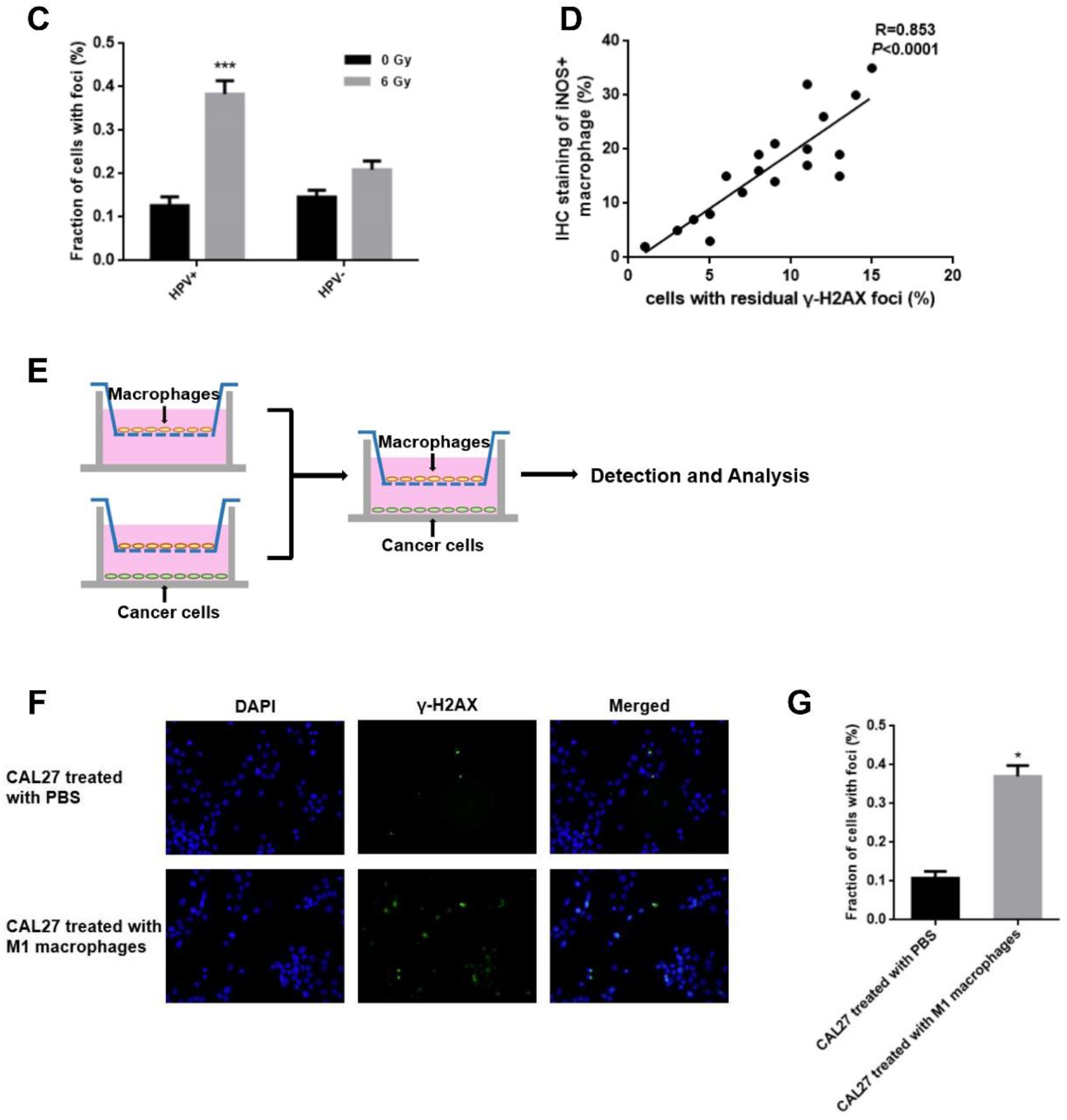
M1 macrophage polarization induced by HPV+ HNSCC-derived exosomes increase tumor radiosensitivity. **A**, IHC analysis of iNOS and CD163 expression in HNSCCs. **B**, Immunofluoresence analysis of γ-H2AX foci expression in HNSCCs 24h after irradiation. **C**, Quantitation of γ-H2AX foci in HNSCCs 24h after irradiation. **D**, Correlation analysis between the fraction of γ-H2AX foci and IHC score of iNOS+ macrophages. **E**, Schematic illustration of the *in vitro* indirect co-culture system. **F**, Immunofluorescence detection of γ-H2AX foci in CAL27 cells 24h after co-culture with M1 macrophages. **G**, Quantitation of γ-H2AX foci in CAL27 cells 24h after irradiation. \**P*<0.05, \*\**P*<0.01, \*\*\**P*<0.001.

We used an *in vitro* indirect co-culture system to study the role of M1 macrophages in radiosensitivity (Figure 2E). M1 macrophages were co-cultured with HPV-tumor cells CAL27. Then, radiosensitivity of CAL27 cells was measured. Immunofluorescence assay showed that M1 macrophages significantly increased the radiosensitivity of CAL27 cells (*P*<0.05) (Figure 2F-G).

### miR-9 is highly expressed in exosomes derived from HPV+ HNSCC cells and can be transferred to macrophages through exosomes

It has been reported that exosomes contain bioactive molecules, such as miRNAs, which are involved in intercellular communication. Further, exosomal miRNA profiles resembled those of the parent cells. We analyzed the differentially expressed miRNAs (DEmiRNAs; FC≥2, *P*<0.05) between HPV+ and HPV-HNSCCs from TCGA database. The data suggested that miR-9 was highly expressed in HPV+ HNSCCs (Figure 3A). Consistent with TCGA result, miR-9 was also highly expressed in HPV+ HNSCC cells compared with HPV-HNSCC cells (Figure 3B). We then used qPCR and confirmed the expression of miR-9 in exosomes derived from HNSCC cells. The results showed that miR-9 was significantly enriched in HPV+ HNSCC-derived exosomes compared with HPV-group (Figure 3C). RNA sequencing was performed using the exosomal RNA from HNSCC cells (CAL27, SCC47 and SCC90) to confirm whether miR-9 was highly expressed in HPV+ exosomes. As shown in Figure 3D, the expression of miR-9 was markedly up-regulated in HPV+ HNSCC-derived exosomes (SCC47 and SCC90) compared with HPV-HNSCC-derived exosomes (CAL27). Since exosomes can deliver bioactive molecules to other cells, we used macrophages incubated with HNSCC-derived exosomes to detect whether miR-9 was delivered these to macrophages. The data showed that macrophages treated with HPV+ HNSCC-derived exosomes expressed higher miR-9 than HPV-HNSCC-derived exosomes or PBS (Figure 3E). Thus, miR-9 is highly expressed in HPV+ HNSCC-derived exosomes, and can be transferred to human macrophages through exosomes.

**Figure 3.**
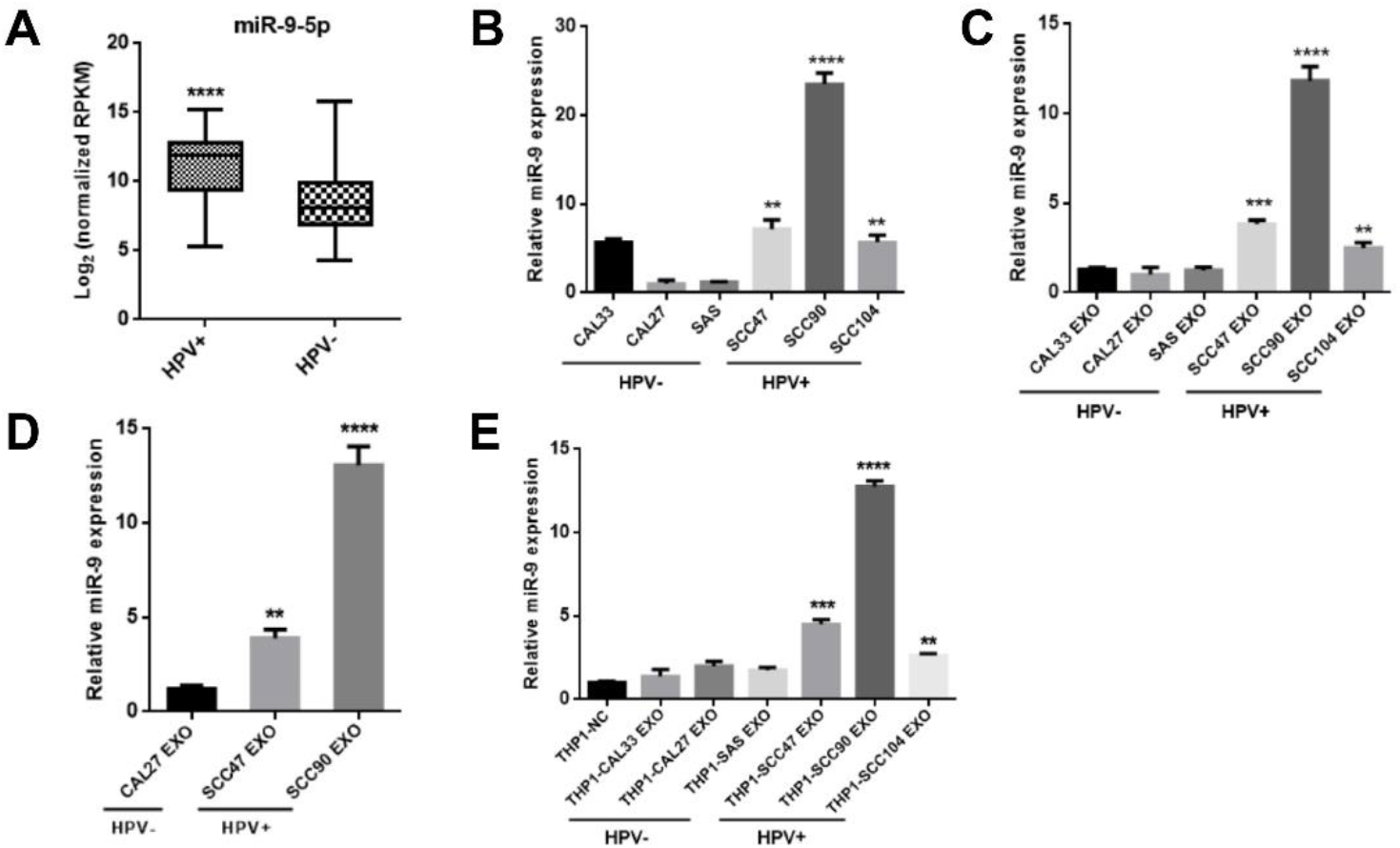
MiR-9 is highly expressed in exosomes derived from HPV+ HNSCC cells and can be transferred to human macrophages through exosomes. **A**, Expression of miR-9 in primary TCGA HNSCCs. **B and C**, Expression of miR-9 was analysed by qPCR in HNSCC cells and their derived exosomes. **D**, Expression of miR-9 in exosomes as detected by RNA sequencing. **E**, Macrophages were cultured with HNSCC-derived exosomes for 48h. miR-9 expression levels were examined in these macrophages by qPCR. \**P*<0.05, \*\**P*<0.01, \*\*\**P*<0.001, \*\*\*\**P*<0.0001.

### Exosomal miR-9 induces the polarization of M1 macrophages to increase the radiosensitivity of HNSCCs via down-regulation of PPARδ expression

To demonstrate the function of miR-9 in the polarization of M1 phenotype macrophages, we employed miR-9 mimic to modulate miR-9 levels in macrophages. M1 polarized macrophages induced by treatment of THP1 cells with LPS and IFNγ served as the positive control. IHC results showed that miR-9 significantly up-regulated expression of M1 markers (iNOS and TNF-α) (Figure 4A). ELISA assay showed that LPS+IFNγ treatment induced the secretion of IL-6 and TNFα, but reduced the secretion of IL-10. miR-9 significantly up-regulated the expression of IL-6 and TNFα, and down-regulated the expression of IL-10 in THP1 cells (Figure 4B-D). Furthermore, we examined whether miR-9-overexpressing macrophages increased the radiosensitivity of HNSCC cells. We found that the radiosensitivity of HPV-HNSCC cells (CAL27) was significantly increased by macrophages transfected with miR-9 mimics (*P*<0.01) (Figure 4E).

**Figure 4.**
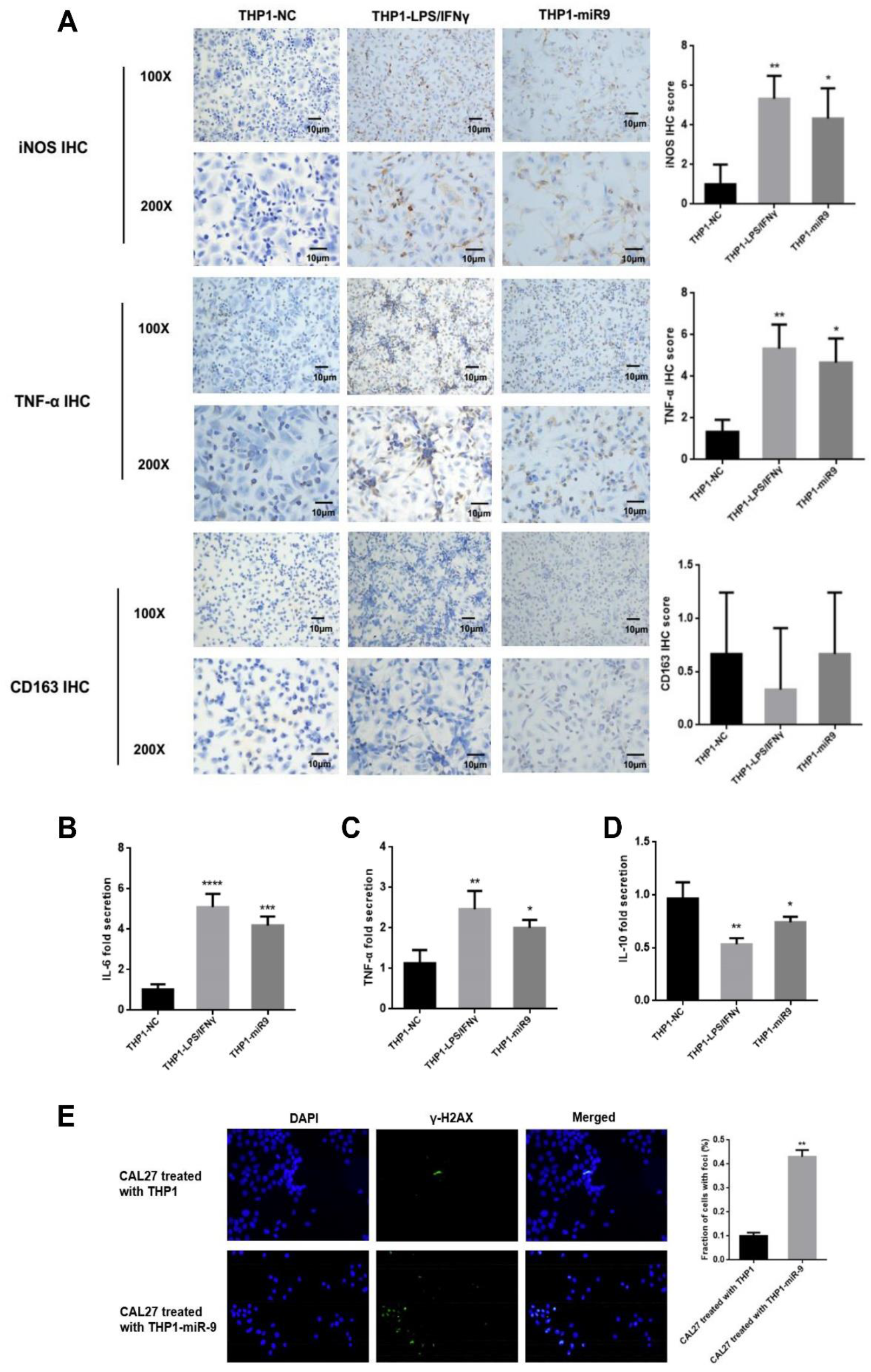
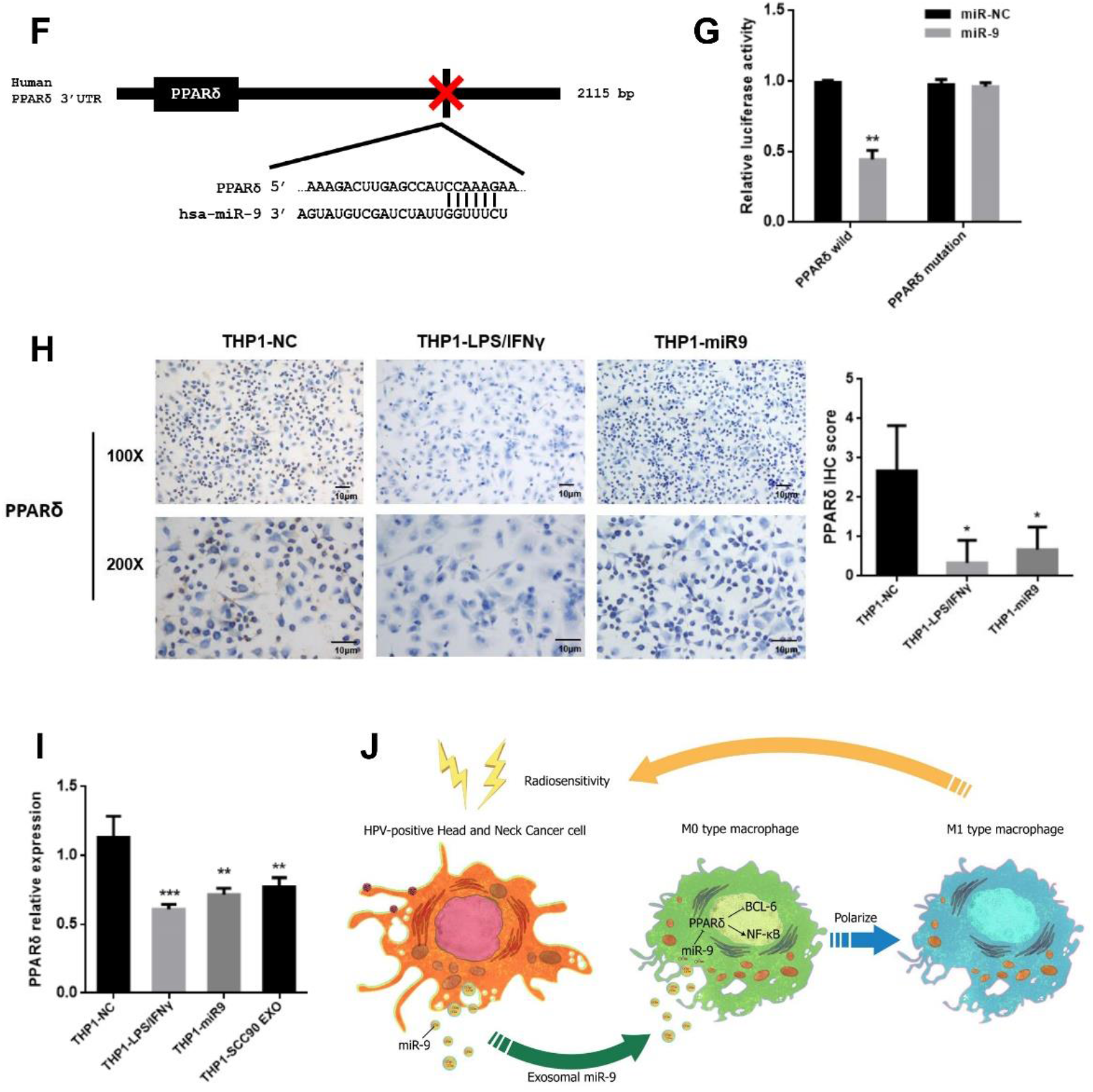
Exosomal miR-9 induces the polarization of M1 macrophages to increase the radiosensitivity of HNSCC. **A**, IHC analysis of iNOS, TNF-α and CD163 expression in THP1-derived macrophages treated with miR-9 mimic. **B,C and D**, Cytokines levels in miR-9 mimic-treated THP1. **E**, Immunofluorescence-based detection of γ-H2AX foci 24h after irradiation of CAL27 cells. **F**, Binding site of miR-9-5p in the 3’UTR of the PPARδ gene, according to bioinformatic analysis. **G**, 293T cells were co-transfected with the 3′UTR luciferase reporter plasmid (500ng) and miR-9-5p mimics or miR-negative control (50nM). Luciferase activity was measured after 40h. **H**, IHC analysis of PPARδ inTHP1-derived macrophages treated with miR-9 mimic. **I**, qPCR analysis of PPARδ expression in THP1-derived macrophages treated with miR-9 mimic. **J**, Schematic model of HPV+ HNSCC-derived exosomal miR-9 promoting macrophage polarization and increasing radiosensitivity of HNSCC. \**P*<0.05, \*\**P*<0.01, \*\*\**P*<0.001, \*\*\*\**P*<0.0001.

To explore the mechanisms underlying exosomal miR-9-mediated induction of macrophage polarization, we predicted its target genes by TargetScan. PPARδ was identified as a potential target (Figure 4F). Luciferase assay was performed to confirm target-miRNA interaction. Our data showed that miR-9 decreased the luciferase activity of Luc-PPARδ-3’ UTR, but had a minimal effect on the negative control (Figure 4G). IHC results showed that LPS+IFNγ down-regulated the expression of PPARδ, while overexpression of miR-9 significantly inhibited the expression of PPARδ in macrophages (Figure 4H). We used the exosomes derived from HPV+ HNSCC cells to treat macrophages and found that HPV+ HNSCC-derived exosomes significantly inhibited PPARδ mRNA levels in macrophages (*P*<0.01) (Figure 4I).

## Discussion

HPV is an important risk factor for HNSCC, and is associated with tumor biology and clinical characteristics including response to radiation(3,6,20). Recently, the involvement of HPV in the radiosensitivity of cancers has been established through altered tumor immune microenvironment. Exosomes released from cancer cells can induce alterations in the tumor microenvironment (21–25). In our study, we isolated exosomes from culture medium of HNSCC cells and found that HPV promoted the release of exosomes from HNSCC cells. Overexpression of exosomal miR-9 induced the polarization of M1 macrophages and then enhanced the response of tumor cells to radiation. This suggests that miR-9 may be a potential treatment strategy for HNSCC.

Exosomes are lipid-bilayer-enclosed extracellular vesicles that can be released from different cell types, especially tumor cells (26,27). Increasing evidence strongly suggests that exosomes can promote activation of immune response in the tumor microenvironment by communicating with tumor cells (28,29). Tumor-derived exosomes activated CD4+ T cells to promote the mitochondrial apoptotic pathway (30). Macrophages are a major component of the tumor-infiltrating immune cell population. The tumor-associated macrophages can be divided into types – M1 and M2 – that function as pro-inflammatory and anti-inflammatory agents, respectively. M1 macrophages produce iNOS, which enhances tumor radiosensitivity (31,32). In this study, we found that HPV+ HNSCC cells released exosomes that induced the polarization of macrophages to the M1 phenotype, thereby increasing tumor response to radiation *in vitro*. Our results suggest that HPV+ HNSCC-derived exosomes are important components of the tumor microenvironment and have an important contribution to the crosstalk between tumor cells and immune cells.

miRNAs are small non-coding RNAs that can downregulate genes by binding to the 3′untranslated-region (UTR) of specific target mRNAs. Valadi et al. proved that miRNAs could be isolated from host cells and transferred to the surrounding target cells through exosomes (33). miR-9 is a functional miRNA that possesses functions crucial for cancer growth, metastasis, immunity, and radiosensitivity (34–36). Previous studies have reported that HPV-mediated activation by miR-9 in both cervical cancer and HNSCC (37–39). In this study, we demonstrated that miR-9 was significantly enriched in exosomes secreted from HPV+ HNSCC cells and could be transferred to macrophages via exosomes. HPV increased the expression of miR-9 in both HNSCC cells and HNSCC cells-dervied exosomes. miR-9 overexpression in HNSCC cells increased miR-9 levels in the secreted exosomes. This promoted the polarization of M1 macrophages, which was followed by enhanced radiosensitivity of HNSCC cells *in vitro*. These results suggest that HPV+ HNSCC cells-derived exosomal miR-9 play an important role in the tumor immune microenvironment and tumor response to radiation.

PPARδ is a ubiquitously expressed member of the ligand-activated nuclear receptor superfamily and plays an important role in inflammation and innate immunity. Upon ligand binding, PPARδ gets activated and releases the anti-inflammatory transcriptional suppressor BCL-6, which leads to suppression of inflammatory gene expression. PPARδ has been shown to drive transcriptional programming of M1 phenotype macrophages (35,40). M1 macrophages expressed higher levels of iNOS, which resulted in higher levels of NO and enhanced radiosensitivity of tumor cells (41). In our study, we showed that downregulation of PPARδ by exosomal miR-9 promoted the polarization of M1 macrophages and increased the expression of iNOS. In addition, M1 macrophages enhanced the sensitization of HNSCC cells to radiation.

In summary, we demonstrated that exosomal miR-9 overexpression promoted an improved response to radiation in HNSCC. HPV promoted the secretion of exosomal miR-9 from HNSCCs. miR-9-rich exosomes induced the polarization of M1 macrophages via downregulation of PPARδ, thereby resulting in enhanced sensitivity of HNSCCs to radiation (Figure 4J). Our findings demonstrated the role of exosomal miR-9 in radiosensitivity of HNSCC and may provide a potential treatment strategy for HNSCC.

## Acknowledgments

We thank Ji Sun from Harbin Medical University Cancer Hospital for providing clinical tumor samples. This study was supported by National Natural Science Foundation of China [grant number 81672670]; Graduate innovation research project of Harbin Medical University [grant number YJSCX2017-8HYD] and Heilongjiang Provincial Science and Technology Innovation Team in Higher Education Institute for Infection and Immunity.

